# 3D Ground Truth Annotations of Nuclei in 3D Microscopy Volumes

**DOI:** 10.1101/2022.09.26.509542

**Authors:** Alain Chen, Liming Wu, Seth Winfree, Kenneth W. Dunn, Paul Salama, Edward J. Delp

**Affiliations:** Video and Image Processing Laboratory, School of Electrical and Computer Engineering, Purdue University, West Lafayette, Indiana; Pathology and Microbiology Department, University of Nebraska Medical Center, Omaha, Nebraska; Division of Nephrology, School of Medicine, Indiana University, Indianapolis, Indiana; Department of Electrical and Computer Engineering, Indiana University-Purdue University, Indianapolis, Indiana

**Keywords:** ground truth, segmentation, training data

## Abstract

In this paper we describe a set of 3D microscopy volumes we have partially manually annotated. We describe the volumes annotated and the tools and processes we use to annotate the volumes. In addition, we provide examples of annotated subvolumes. We also provide synthetically generated 3D microscopy volumes that can be used for training segmentation methods. The full set of annotations, synthetically generated volumes, and original volumes can be accessed as described in the paper.

## 1. INTRODUCTION

Recent progress in microscopy technology has enabled the acquisition of large 3D volumetric data, including 3D multi-spectral data, using fluorescence imaging. Analyses of these large volumes often involve a segmentation of cells or cellular nuclei. As cell boundaries are typically poorly defined in tissues, segmentation can be done on nuclei rather than entire cells [1]. One then characterize and classify cells on the basis of the fluorescence in the user-defined regions surrounding the nuclei [2].

Segmentation techniques based on deep learning have shown great promise, in some cases providing accurate and robust results across a range of image types [3, 4, 5, 6, 7]. However, their utility is limited by the large amounts of manually annotated data needed for training, validation, and testing. Annotation is a labor-intensive and time-consuming process, especially for 3D volumes.

Many publicly available annotated microscopy datasets, such as those in the Broad Bioimage Benchmark Collection [8], are 2D images. There is a lack of 3D volumes with annotations that are available. The 3D volumes that are available are typically synthetic volumes and not real microscopy volumes.

In this paper we describe a set of annotated 3D subvolumes of real microscopy volumes, along with a set of synthetically generated subvolumes, that are available for use by the research community. A list of available volumes can be found in Tables 1, 2, and 3.

**Table 1.** Table of Volumes. Subvolumes can be found in https:doi.org//10.5281/zenodo.7065147

| Index | Volume Name | Original Volume Repository | Number of Subvolumes Annotated | Number of Synthetic Subvolumes | Synthetic Subvolume Directory Name |
| --- | --- | --- | --- | --- | --- |
| 1 | *Cleared_mouse_intestine_1* | https://doi.org/10.5281/zenodo.7032409 | 4 | 4 | Cleared_mouse_intestine_1/synthetic |
| 2 | *Diabetic_Biopsy_Human_Spectral_1* | https://doi.org/10.5281/zenodo.7023876 | 5 | 50 | Diabetic_Biopsy_Human_Spectral_1/synthetic |
| 3 | *Diabetic_Biopsy_Human_Spectral_3* | https://doi.org/10.5281/zenodo.7023871 | 6 | 50 | Diabetic_Biopsy_Human_Spectral_3/synthetic |
| 4 | *Kidney_Cortex_Human_Spectral_1* | https://doi.org/10.5281/zenodo.5842278 | 6 | 92 | Kidney_Cortex_Human_Spectral_1/synthetic |
| 5 | *BABB-cleared_kidney_1* | https://doi.org/10.5281/zenodo.7032413 | 4 | 50 | BABB-cleared_kidney_1/synthetic |
| 6 | *Kidney_Human_Nephrectomy_1* | https://doi.org/10.5281/zenodo.7023880 | 4 | 4 | Kidney_Human_Nephrectomy_1/synthetic |
| 7 | *Scale-cleared_rat_kidney_1* | https://doi.org/10.5281/zenodo.7032415 | 1 | 50 | Scale-cleared_rat_kidney_1/synthetic |
| 8 | *Rat_liver_1* | https://doi.org/10.5281/zenodo.7032419 | The entire volume is annotated. | 50 | Rat_liver_1/synthetic |

**Table 2.** Table of Annotated Subvolumes. Files can be found in https:doi.org//10.5281/zenodo.7065147

| Index | Volume Name | Subvolume File Names | Annotated File Names |
| --- | --- | --- | --- |
| 1 | ***Cleared mouse intestine 1*** | Cleared_mouse_intestine_1/<br>Cleared_mouse_intestine_1_150_800_20/<br>Cleared_mouse_intestine_1_150_800_20.tif<br>Cleared_mouse_intestine_1/<br>Cleared_mouse_intestine_1_400_300_120/<br>Cleared_mouse_intestine_1_400_300_120.tif<br>Cleared_mouse_intestine_1/<br>Cleared_mouse_intestine_1_400_850_57/<br>Cleared_mouse_intestine_1_400_850_57.tif<br>Cleared_mouse_intestine_1/<br>Cleared_mouse_intestine_1_430_315_74/<br>Cleared_mouse_intestine_1_430_315_74.tif | Cleared_mouse_intestine_1/<br>Cleared_mouse_intestine_1_150_800_20/<br>Cleared_mouse_intestine_1_150_800_20_gt.tif<br>Cleared_mouse_intestine_1/<br>Cleared_mouse_intestine_1_400_300_120/<br>Cleared_mouse_intestine_1_400_300_120_gt.tif<br>Cleared_mouse_intestine_1/<br>Cleared_mouse_intestine_1_400_850_57/<br>Cleared_mouse_intestine_1_400_850_57_gt.tif<br>Cleared_mouse_intestine_1/<br>Cleared_mouse_intestine_1_430_315_74/<br>Cleared_mouse_intestine_1_430_315_74_gt.tif |
| 2 | ***Diabetic Biopsy Human Spectral 1*** | Diabetic_Biopsy_Human_Spectral_1/Diabetic<br>_Biopsy_Human_Spectral_1-3100-1700/Diabetic<br>_Biopsy_Human_Spectral_1-3100-1700.tif<br>Diabetic_Biopsy_Human_Spectral_1/Diabetic<br>_Biopsy_Human_Spectral_1-5300-2300/Diabetic<br>_Biopsy_Human_Spectral_1-5300-2300.tif<br>Diabetic_Biopsy_Human_Spectral_1/Diabetic<br>_Biopsy_Human_Spectral_1-5800-2500/Diabetic<br>_Biopsy_Human_Spectral_1-5800-2500.tif<br>Diabetic_Biopsy_Human_Spectral_1/Diabetic<br>_Biopsy_Human_Spectral_1-6000-1150/Diabetic<br>_Biopsy_Human_Spectral_1-6000-1150.tif<br>Diabetic_Biopsy_Human_Spectral_1/Diabetic<br>_Biopsy_Human_Spectral_1-8000-1350/Diabetic<br>_Biopsy_Human_Spectral_1-8000-1350.tif | Diabetic_Biopsy_Human_Spectral_1/Diabetic<br>_Biopsy_Human_Spectral_1-3100-1700/Diabetic<br>_Biopsy_Human_Spectral_1-3100-1700_gt.tif<br>Diabetic_Biopsy_Human_Spectral_1/Diabetic<br>_Biopsy_Human_Spectral_1-5300-2300/Diabetic<br>_Biopsy_Human_Spectral_1-5300-2300_gt.tif<br>Diabetic_Biopsy_Human_Spectral_1/Diabetic<br>_Biopsy_Human_Spectral_1-5800-2500/Diabetic<br>_Biopsy_Human_Spectral_1-5800-2500_gt.tif<br>Diabetic_Biopsy_Human_Spectral_1/Diabetic<br>_Biopsy_Human_Spectral_1-6000-1150/Diabetic<br>_Biopsy_Human_Spectral_1-6000-1150_gt.tif<br>Diabetic_Biopsy_Human_Spectral_1/Diabetic<br>_Biopsy_Human_Spectral_1-8000-1350/Diabetic<br>_Biopsy_Human_Spectral_1-8000-1350_gt.tif |
| 3 | ***Diabetic Biopsy Human Spectral 3*** | Diabetic_Biopsy_Human_Spectral_3/Diabetic<br>_Biopsy_Human_Spectral_3-1529_864/Diabetic<br>_Biopsy_Human_Spectral_3-1529_864.tif<br>Diabetic_Biopsy_Human_Spectral_3/Diabetic<br>_Biopsy_Human_Spectral_3-1728_1100/Diabetic<br>_Biopsy_Human_Spectral_3-1728_1100.tif<br>Diabetic_Biopsy_Human_Spectral_3/Diabetic<br>_Biopsy_Human_Spectral_3-3480_1070/Diabetic<br>_Biopsy_Human_Spectral_3-3480_1070.tif<br>Diabetic_Biopsy_Human_Spectral_3/Diabetic<br>_Biopsy_Human_Spectral_3-4180_1590/Diabetic<br>_Biopsy_Human_Spectral_3-4180_1590.tif<br>Diabetic_Biopsy_Human_Spectral_3/Diabetic<br>_Biopsy_Human_Spectral_3-6137_1985/Diabetic<br>_Biopsy_Human_Spectral_3-6137_1985.tif<br>Diabetic_Biopsy_Human_Spectral_3/Diabetic<br>_Biopsy_Human_Spectral_3-6320_1385/Diabetic<br>_Biopsy_Human_Spectral_3-6320_1385.tif | Diabetic_Biopsy_Human_Spectral_3/Diabetic<br>_Biopsy_Human_Spectral_3-1529_864/Diabetic<br>_Biopsy_Human_Spectral_3-1529_864_gt.tif<br>Diabetic_Biopsy_Human_Spectral_3/Diabetic<br>_Biopsy_Human_Spectral_3-1728_1100/Diabetic<br>_Biopsy_Human_Spectral_3-1728_1100_gt.tif<br>Diabetic_Biopsy_Human_Spectral_3/Diabetic<br>_Biopsy_Human_Spectral_3-3480_1070/Diabetic<br>_Biopsy_Human_Spectral_3-3480_1070_gt.tif<br>Diabetic_Biopsy_Human_Spectral_3/Diabetic<br>_Biopsy_Human_Spectral_3-4180_1590/Diabetic<br>_Biopsy_Human_Spectral_3-4180_1590_gt.tif<br>Diabetic_Biopsy_Human_Spectral_3/Diabetic<br>_Biopsy_Human_Spectral_3-6137_1985/Diabetic<br>_Biopsy_Human_Spectral_3-6137_1985_gt.tif<br>Diabetic_Biopsy_Human_Spectral_3/Diabetic<br>_Biopsy_Human_Spectral_3-6320_1385/Diabetic<br>_Biopsy_Human_Spectral_3-6320_1385_gt.tif |

**Table 3.**
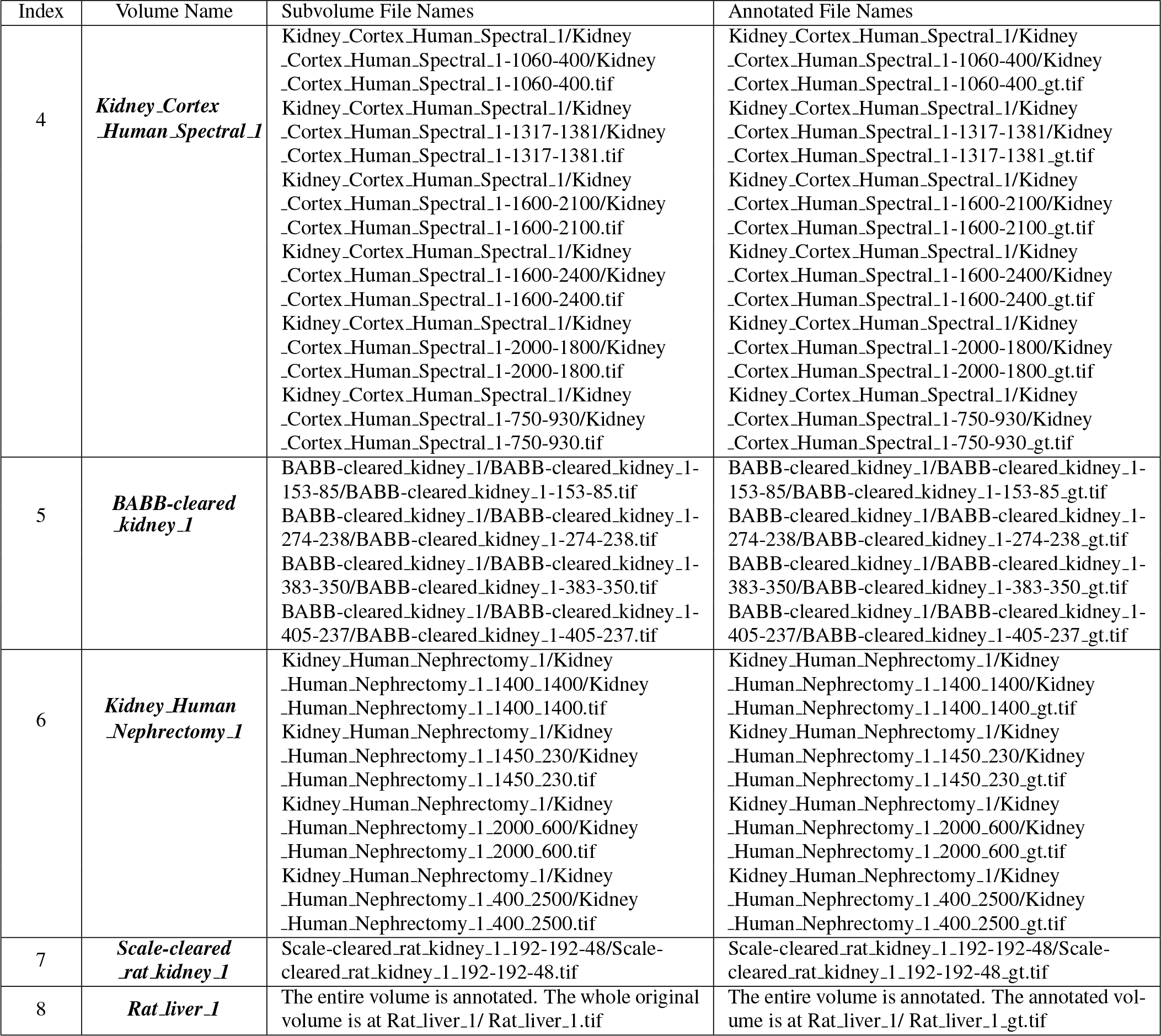
Table of Annotated Subvolumes Continued. Files can be found in https:doi.org//10.5281/zenodo.7065147

## 2. MANUAL ANNOTATION PROCESS

In this section, we describe the process we used to annotate real 3D microscopy volumes.

Thirty 3D subvolumes (of 128 by 128 pixels by variable depth in *z*) from eight different 3D microscopy volumes were annotated. Note that we manually annotated the nuclei in the subvolumes. Sub-volumes from each volume were chosen such that the subvolumes are different and representative areas of the larger original volume. Subvolumes were also chosen from particular regions of interest, such as regions with overlapping, densely-packed, or irregularly shaped nuclei.

### 2.1. Annotating Using ITK-Snap

Manual annotations were obtained using ITK-Snap [9]. Here we describe the specific steps in ITK-Snap we use to annotate the sub-volumes.

We use the 3D adaptive brush as a first pass for annotating a subvolume. Different “Active Labels” are used for nuclei that touch other nuclei. It is acceptable for nuclei that are far away (i.e. not touching) to have the same “Active Label.” As a result of this, as many as eight different “Active Labels” are used for each subvolume. Minor adjustments were made using a 3D round brush after the initial use of the 3D adaptive brush.

The final result is a manually annotated subvolume with the same size as the original subvolume. Any background voxel, i.e. any voxel not being annotated as part of a nucleus, has a voxel intensity of 0. A voxel belonging to a nucleus will have an intensity value between 1 and *m*, where *m* is the number of “Active Labels” used during the annotation. Each intensity value between 1 and *m* corresponds to an “Active Label.”

## 3. SYNTHETICALLY GENERATED VOLUMES

We also generate synthetic microscopy subvolumes using SpCycleGAN [10]. In SpCycleGAN, we generate synthetic nuclei segmentation masks where we can control the size, location, shape, and orientation of desired nuclei. SpCycleGAN learns the properties of real microscopy images and then generates synthetic microscopy images of nuclei with the size, location, shape, and orientation of the nuclei in the synthetic nuclei segmentation masks. Note that only microscopy volumes and synthetically generated nuclei segmentation masks are input to the subvolume generation process. Manual annotations were not needed for synthetic microscopy subvolume generation because we know the location of the nuclei as part of the generation process. The synthetic volumes in Table 1 were generated using SpCycleGAN as described in [10, 6, 11]. Please refer to these publications [10, 6, 11] for a more detailed description of SpCycleGAN and examples of how synthetic subvolumes were generated.

## 4. DESCRIPTION OF ANNOTATED SUBVOLUMES

We obtained manually annotated subvolumes from eight different microscopy volumes. Here we describe each of the microscopy volumes in detail.

1. The ***Cleared mouse intestine 1*** volume is a 512 × 930 1× 57 (*X* × *Y* × *Z*) volume of cleared mouse intestine tissue. The voxel dimensions are 1 × 1 × 1 micron^3^ (*X* × *Y* × *Z*). Images of paraformaldehyde-fixed mouse intestine were labeled with DAPI and imaged using confocal microscopy with a Leica SP8 confocal/multiphoton microscope using a 20X NA 0.75 multi-immersion objective. Tissues were cleared using a modified version of previously described procedures [12].
2. The ***Diabetic Biopsy Human Spectral 1*** volume is a 9464 × 2877 × 35 (*X* × *Y* × *Z*) volume of diabetic kidney biopsy tissue. The voxel dimensions are 0.5407 × 0.5408 × 1.0412 micron^3^ (*X* × *Y* × *Z*). Fresh-frozen human kidney samples are placed in cold Optimal Cutting Temperature (OCT) compound for 3 min and then transferred to a cryomold with partially frozen OCT in the bottom on a block of dry ice. Once the OCT is completely frozen, the tissue block is wrapped in parafilm and stored at 80 ^*◦*^ C. Frozen tissues are sectioned to a thickness of 50 µm using a cryostat and then immediately fixed in 4% fresh paraformaldehyde (PFA) for 24 h, and subsequently stored at 4 ^*◦*^ C in 0.25% PFA. Tissue was imaged using a Leica SP8 confocal scan-head mounted to an upright DM6000 microscope with computer-controlled motorized stage [13].
3. The ***Diabetic Biopsy Human Spectral 3*** volume is a 12160 × 2440 × 36 (*X* × *Y* × *Z*) volume of diabetic kidney biopsy tissue. The voxel dimensions are 0.5406 × 0.5408 × 1.0412 micron^3^ (*X* × *Y* × *Z*). Fresh-frozen human kidney samples are placed in cold Optimal Cutting Temperature (OCT) compound for 3 min and then transferred to a cryomold with partially frozen OCT in the bottom on a block of dry ice. Once the OCT is completely frozen, the tissue block is wrapped in parafilm and stored at 80 ^*◦*^ C. Frozen tissues are sectioned to a thickness of 50 µm using a cryostat and then immediately fixed in 4% fresh paraformaldehyde (PFA) for 24 h, and subsequently stored at 4 ^*◦*^ C in 0.25% PFA. Tissue was imaged using a Leica SP8 confocal scan-head mounted to an upright DM6000 microscope with computer-controlled motorized stage [13].
4. The ***Kidney Cortex Human Spectral 1*** volume is a 3889 × 4734 × 19 (*X* × *Y* × *Z*) volume of human nephrectomy tissue. The voxel dimensions are 0.5407 × 0.5407 × 1.0412 micron^3^ (*X* × *Y* × *Z*). Tissue was cryosectioned and fixed in formaldehyde (4.0% and stored in 0.25% in 1X PBS) and imaged using an FV1000 (Olympus) microscope at 20x 0.75 NA oil immersion objective [14].
5. The ***BABB-cleared kidney 1*** volume is a 512 × 512 × 415 (*X* × *Y* × *Z*) volume of BABB-cleared rat kidney. The voxel dimensions are 1 × 1 × 1 micron^3^ (*X* × *Y* × *Z*). Images of paraformaldehyde-fixed rat kidney tissue were collected with a 40X NA 1.3 oil immersion objective, using an Olympus FV1000 confocal microscope system (Olympus America, Inc., Center Valley, PA, USA) adapted for two-photon microscopy. Rat kidney tissues were fixed, cleared and imaged using confocal microscopy (anti-vimentin immunofluorescence, and Lens culinaris agglutinin) and multiphoton microscopy (Hoechst33342-labeled nuclei) as previously described [15].
6. The ***Kidney Human Nephrectomy 1*** volume is a 2912 × 3520 × 35 (*X* × *Y* × *Z*) volume of human stone disease biopsy. Tissue was fixed in 4% paraformaldehyde overnight and stored in 0.25% paraformaldehyde. Nuclei were stained with DAPI and imaged by fluorescence confocal imaging microscopy [16]. The voxel dimensions are 1 *×* 1 *×* 1 micron^3^ (*X × Y × Z*).
7. The ***Scale-cleared rat kidney 1*** volume is a 512 *×* 512 *×* 200 (*X × Y × Z*) volume of scale-cleared rat kidney. The voxel dimensions are 1 *×* 1 *×* 1 micron^3^ (*X × Y × Z*). An Olympus Fluoview 1000 MPE confocal/multiphoton microscope system mounted on an Olympus IX-81 inverted stand (Olympus America, Inc., Center Valley, PA, USA), equipped with an Olympus 60X oil immersion objective was used to collect images of rat kidney. For this volume, paraformaldehyde-fixed tissue was labeled with phalloidin and Hoechst 33342, cleared and mounted in Scale mounting medium [17] and imaged by confocal microscopy using an Olympus 25X, NA1.05 water immersion objective.
8. The ***Rat liver 1*** volume is a 512 *×* 512 *×* 32 (*X × Y × Z*) volume of rat liver. The voxel dimensions are 1 *×* 1 *×* 1 micron^3^ (*X Y Z*). An Olympus Fluoview 1000 MPE confocal/multiphoton microscope system mounted on an Olympus IX-81 inverted stand (Olympus America, Inc., Center Valley, PA, USA), equipped with an Olympus 60X oil immersion objective was used to collect images of rat liver. For this volume, paraformaldehyde-fixed rat liver tissue was labelled with phalloidin, anti-Mrp2 immunofluorescence, and Hoechst 33342, cleared and mounted in Scale mounting medium [17] and imaged by confocal microscopy using an Olympus 25X, NA1.05 water immersion objective.

Here we also describe the naming convention of the subvolumes and annotations of the data provided. The main directory contains eight directories, one for each of the volumes provided. Within each of these directories will contain many subdirectories. Each subdirectory corresponds to a subvolume of the original microscopy volume. The subdirectory name will contain the original microscopy volume name and the location of the subvolume in the original volume. Each subdirectory will contain two files: one file corresponding to the real microscopy subvolume, and one file corresponding to the manually annotated subvolume. The annotated file name will contain the suffix ‘ gt’ before the file extension. Under each main directory for each volume, there is a directory named ‘synthetic’ where synthetically generated subvolumes are provided. Under the ‘synthetic’ directory will be two directories, ‘gt’ and ‘syn’. ‘gt’ will contain numbered .tif files containing the masks corresponding to where the nuclei locations will be in the synthetically generated subvolumes. ‘syn’ will contain numbered .tif files containing the synthetically generated microscopy subvolumes. The numbering of the files in ‘gt’ corresponds to the numbering of the files in ‘syn’.

## 5. EXAMPLE GROUND TRUTH ANNOTATIONS

In this section, we give a few examples of the output of our 3D ground truth annotations. Table 4 shows eight contiguous 2D slices from an annotated 3D subvolume of ***BABB-cleared kidney 1***. Note that the same nucleus across different focal planes is given the same “Active Label” and thus has the same pixel intensity shown in the “Slices of Annotated Subvolume” section. Also note that different nuclei that are close to one another are given different “Active Labels” and thus have different pixel intensities. As indicated in Section 2.1, nuclei that are far away may have the same “Active Label.” Table 5 shows example 2D slices from annotated 3D subvolumes of each volume provided.

**Table 4.**
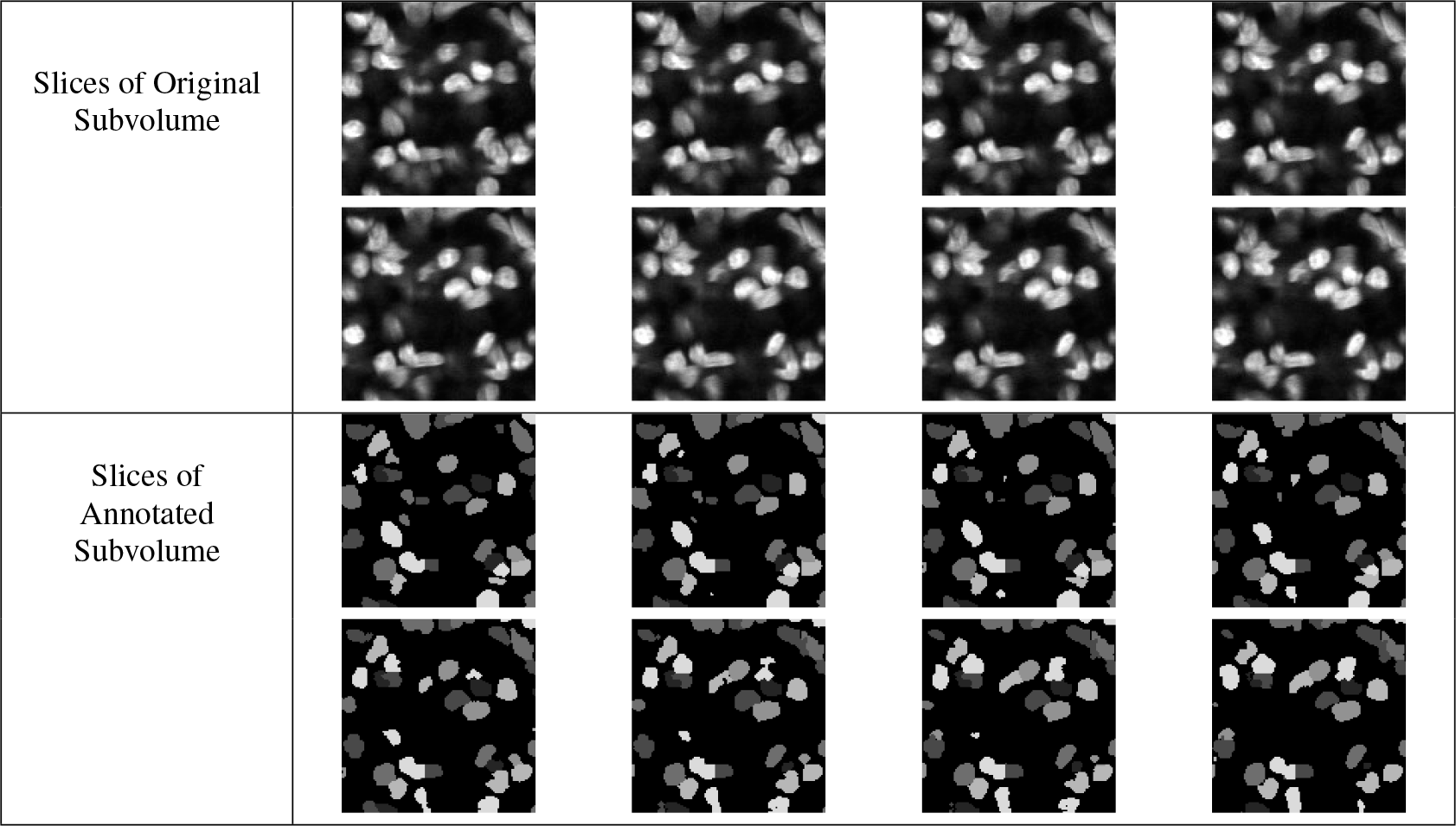
Eight Contiguous 2D Slices from an Annotated 3D Subvolume of the *BA-BB-cleared_kidney_1 Volume*

**Table 5.**
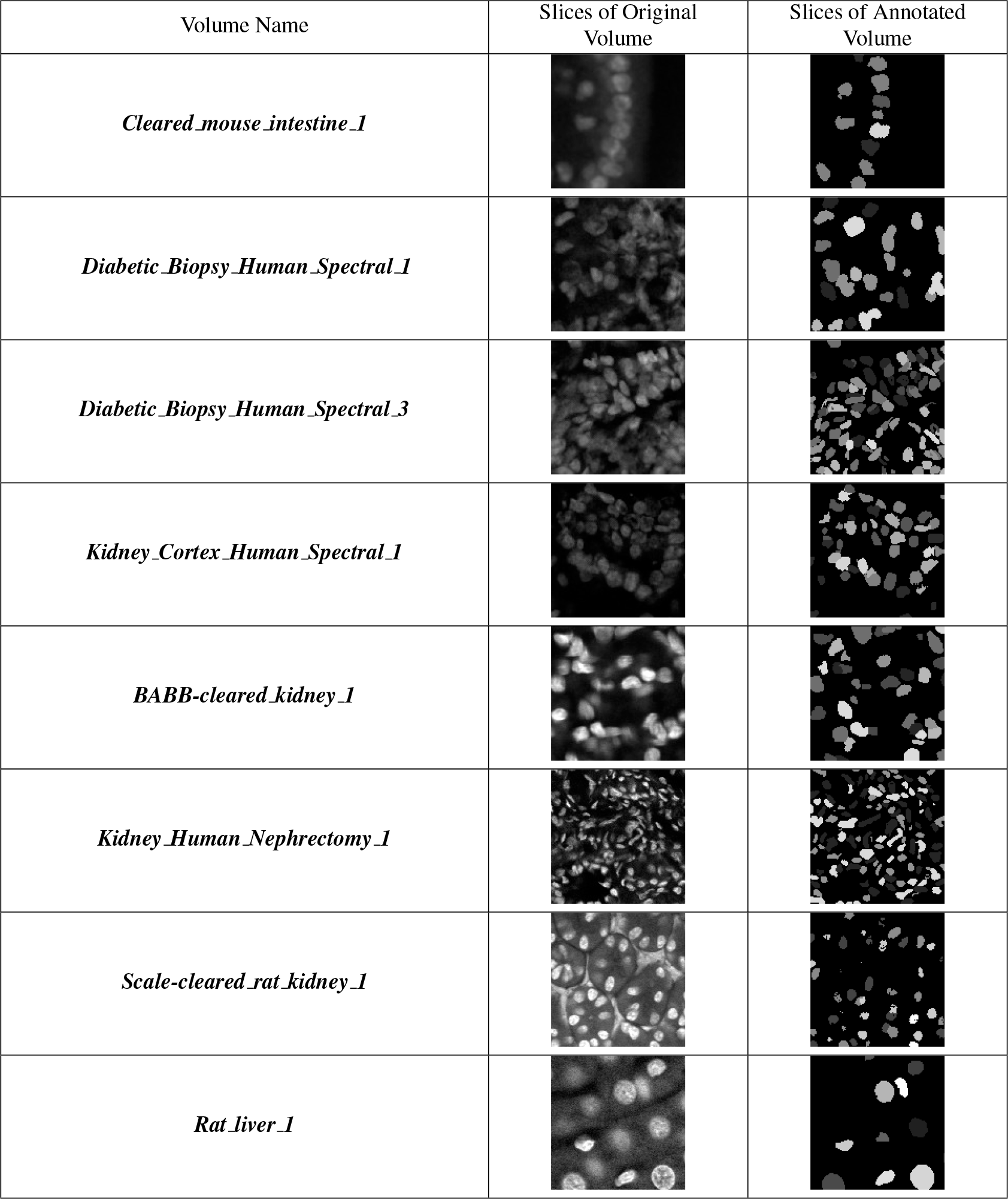
Example 2D Slices from Annotated 3D Subvolumes

### 5.1. Use of Annotated Volumes in Our Work

A subset of these annotated volumes were used to evaluate the quantitative performance of DeepSynth [6], NisNet3D [18], 3D CentroidNet [19], EMR-CNN [20], and RNN-SliceNet [21]. A partial subset of these annotated volumes is also used to lightly retrain a version of NisNet3D [18].

## 6. AVAILABILITY OF DATA

Manually annotated and synthetically generated volumes can be accessed at https:doi.org//10.5281/zenodo.7065147. For a list of file names in the directory, please refer to Table 1. For a list of annotated subvolumes in the directory, please refer to Tables 2 and 3. The synthetically generated and annotated set is 1.67GB. The real, original microscopy volumes are found at their respective repositories according to Table 1. The original volumes amount to 3.13GB.

The data is distributed under Creative Commons license Attribution - NonCommercial - NoDerivs - CC BY-NC-ND.

You are free to:

- Share - copy and redistribute the material in any medium or format
- The licensor cannot revoke these freedoms as long as you follow the license terms

Under the following terms:

- Attribution - You must give appropriate credit, provide a link to the license, and indicate if changes were made. You may do so in any reasonable manner, but not in any way that suggests the licensor endorses you or your use
- NonCommercial - You may not use the material for commercial purposes
- NoDerivatives - If you remix, transform, or build upon the material, you may not distribute the modified material
- No additional restrictions - You may not apply legal terms or technological measures that legally restrict others from doing anything the license permits
- More details are available in the ”readme” files in the images including how we suggest you cite our work and this paper.

## 7. CONCLUSION

In this paper, we provide a set of annotated 3D subvolumes of real microscopy volumes and describe the tools we use to obtain the manual annotations. We also provide a set of synthetically generated microscopy subvolumes, generated from an SpCycleGAN [7], that is available for training segmentation networks.

## 8. ETHICAL STANDARDS AND COMPLIANCE

The collection of the ***Kidney Cortex Human Spectral 1*** volume, the ***Diabetic Biopsy Human Spectral 1*** volume, and the ***Diabetic Biopsy Human Spectral 3*** volume were approved by the Institutional Review Board of Indiana University No. 1906572234.

The collection of ***Kidney Human Nephrectomy 1*** was approved by the Institutional Review Board of Indiana University No. 1010002261.

## 9. ACKNOWLEDGMENTS

This work was partially supported by a George M. O’Brien Award from the National Institutes of Health under grant NIH/NIDDK P30 DK079312 and the endowment of the Charles William Harrison Distinguished Professorship at Purdue University.

Thirteen graduate students, working 105 hours, were involved in obtaining the manual ground truth annotations.

The authors have no conflicts of interest. Address all correspondence to Edward J. Delp.

We would like to thank Michael Ferkowicz and Mervin Yoder for providing the ***Cleared mouse intestine 1*** volume, Sherry Clendenon and Kenneth Dunn for providing the ***BABB-cleared kidney 1*** volume, Malgorzata M. Kamocka and Kenneth Dunn for providing the ***Scale-cleared rat kidney 1*** volume, Sherry Clendenon and Kenneth Dunn for providing the ***Rat liver 1*** volume, Michael Ferkowicz, Seth Winfree, and Tarek El-Achkar for providing the ***Kidney Cortex Human Spectral 1*** volume, Michael Ferkowicz and Tarek El-Achkar for providing ***Diabetic Biopsy Human Spectral 3***, Michael Ferkowicz, Seth Winfree, and Mervin Yoder for providing the ***Diabetic Biopsy Human Spectral 1*** volume, and Seth Winfree and Tarek El-Achkar for providing ***Kidney Human Nephrectomy 1***.

Please contact for any questions about the data.

